# The Distribution of Bacterial Doubling Times in the Wild

**DOI:** 10.1101/214783

**Authors:** Beth Gibson, Daniel Wilson, Edward Feil, Adam Eyre-Walker

**Affiliations:** School of Life Sciences, University of Sussex, Brighton, BN1 9QG; Nuffield Department of Medicine, University of Oxford, John Radcliff Hospital, Oxford, OX3 9DU; The Milner Centre for Evolution, Department of Biology and Biochemistry, University of Bath, Claverton Down, Bath, BA2 7AY

## Abstract

Generation time varies widely across organisms and is an important factor in the life cycle, life history and evolution of organisms. Although the doubling time (DT), has been estimated for many bacteria in the lab, it is nearly impossible to directly measure it in the natural environment. However, an estimate can be obtained by measuring the rate at which bacteria accumulate mutations per year in the wild and the rate at which they mutate per generation in the lab. If we assume the mutation rate per generation is the same in the wild and in the lab, and that all mutations in the wild are neutral, an assumption that we show is not very important, then an estimate of the DT can be obtained by dividing the latter by the former. We estimate the DT for four species of bacteria for which we have both an accumulation and a mutation rate estimate. We also infer the distribution of DTs across all bacteria from the distribution of the accumulation and mutation rates. Both analyses suggest that DTs for bacteria in the wild are substantially greater than those in the lab, that they vary by orders of magnitude between different species of bacteria and that a substantial fraction of bacteria double very slowly in the wild.

## Introduction

The bacterium *Escherichia coli* can divide every 20 minutes in the laboratory under aerobic, nutrient-rich conditions. But how often does it divide in its natural environment in the gut, under anaerobic conditions where it probably spends much of its time close to starvation? And what do we make of a bacterium, such as *Syntrophobacter fumaroxidans*, which only doubles in the lab every 140 hours (1). Does this reflect a slow doubling time in the wild, or our inability to provide the conditions under which it can replicate rapidly?

Estimating the generation time is difficult for most bacteria in their natural environment and very few estimates are available. The doubling time (DT) for intestinal bacteria has been estimated in several mammals by assaying the quantity of bacteria in the gut and faeces. Assuming no cell death Gibbons & Kapsimalis (1967) (2) estimate the DT for all bacteria in the gut to be 48, 17 and 5.8 hours in hamster, guinea pig and mouse respectively. More recently Yang et al. (2008) (3) have shown that the doubling time of *Pseudomonas aeruginosa* is correlated to cellular ribosomal content *in vitro* and have used this to estimate the DT *in vivo* in a cystic fibrosis patient to be between 1.9 and 2.4 hours.

Although there are very few estimates of the generation time in bacteria, this quantity is important for understanding bacterial population dynamics. Here we use an indirect method to estimate the DT that uses two sources of information. First, we can measure the rate at which a bacterial species accumulates mutations in its natural environment through time using temporarily sampled data (4), or concurrent samples from a population with a known date of origin. We refer to this quantity as the accumulation rate, to differentiate it from the mutation rate, and the substitution rate, the rate at which mutations spread through a species to fixation. If we assume that all mutations in the wild are neutral, an assumption that we show to be unimportant, then the accumulation rate is an estimate of the mutation rate per year, *u*_*y*_. Second, we can estimate the rate of mutation per generation, *u*_*g*_, in the lab using a mutation accumulation experiment and whole genome sequencing, or through fluctuation tests. If we assume that the mutation rate per generation is the same in the wild and in the lab, an assumption we discuss further below, then if we divide the accumulation rate per year in the wild by the mutation rate per generation in the lab, we can estimate the number of generations that the bacterium goes through in the wild and hence the doubling time (DT = 8760 x *u*_*g*_ / *u*_*y*_, where 8760 is the number of hours per year).

## Results

The accumulation rate in the wild and the mutation rate in the lab have been estimated for 34 and 12 bacterial species respectively (Tables S1, S2); we only consider mutation rate estimates from mutation accumulation experiments, since estimates from fluctuation tests are subject to substantial sampling error and unknown bias, and we exclude estimates from hypermutable strains. For four species, *Escherichia coli, Pseudomonas aeruginosa, Salmonella enterica and Vibrio cholera,* we have both an accumulation and a mutation rate estimate and hence can estimate the DT. Amongst these four species we find our DT estimates vary from 1.1 hours in *V. cholerae* to 25 hours in *Salmonella enterica* (Table 1). In all cases the estimated DT in the wild is greater than that of the bacterium in the lab. For example, *E. coli* can double every 20 minutes in the lab but we estimate that it only doubles every 15 hours in the wild.

**Table 1.** Doubling time estimates (hours) for those species for which we have both an estimate of the accumulation and mutation rate. Accumulation rate (AR) references – [1] (5). [2] (6, 7). (7) [3] (8–12). [4] (8, 13). Mutation rate (MR) references – [5] (14). [6] (15). [7] (16); [8] (17).

| Species | Accumulation rate per site per year | Mutation rate per site per generation | DT (hr) (SE) | Lab DT (hr) | Ratio | AR Ref | MR Ref |
| --- | --- | --- | --- | --- | --- | --- | --- |
| *Escherichia coli* | $1.44 \times 10^{-7}$ | $2.54 \times 10^{-10}$ | 15 (7.7) | 0.33 | 45 | 1 | 5 |
| *Pseudomonas aeruginosa* | $3.03 \times 10^{-7}$ | $7.92 \times 10^{-11}$ | 2.3 (0.77) | 0.5 | 4.6 | 2 | 6 |
| *Salmonella enterica* | $2.50 \times 10^{-7}$ | $7.00 \times 10^{-10}$ | 25 (7.9) | 0.5 | 50 | 3 | 7 |
| *Vibrio cholerae* | $8.30 \times 10^{-7}$ | $1.07 \times 10^{-10}$ | 1.1 (0.26) | 0.66 | 1.7 | 4 | 8 |

In theory, it might be possible to estimate the DT in those bacteria for which we have either an accumulation or mutation rate estimate, but not both, by finding factors that correlate with either rate and using those factors to predict the rates. Unfortunately, we have been unable to find any factor that correlates sufficiently well to be usefully predictive. It has been suggested that the mutation rate is correlated to genome size in microbes (18) but, the current evidence for this correlation is very weak, and depends upon the estimate from *Mesoplasma florum* (r = -0.83, *p* < 0.001 with *M. forum* and r = -0.45, *p*= 0.161 without *M. flroum*) (19). However, we can use the accumulation and mutation rate estimates to estimate the distribution of DTs across bacteria. We do this by fitting distributions to the accumulation and mutation rate data, using maximum likelihood, and then dividing one distribution by the other. We assume that both variables are log-normally distributed, an assumption which is supported by Q-Q plots with the exception of the mutation rate per generation in *Mesoplasma florum*, which is a clear outlier (Figure 1.). We repeated all our analyses with and without *M. florum*.

**Figure 1.**
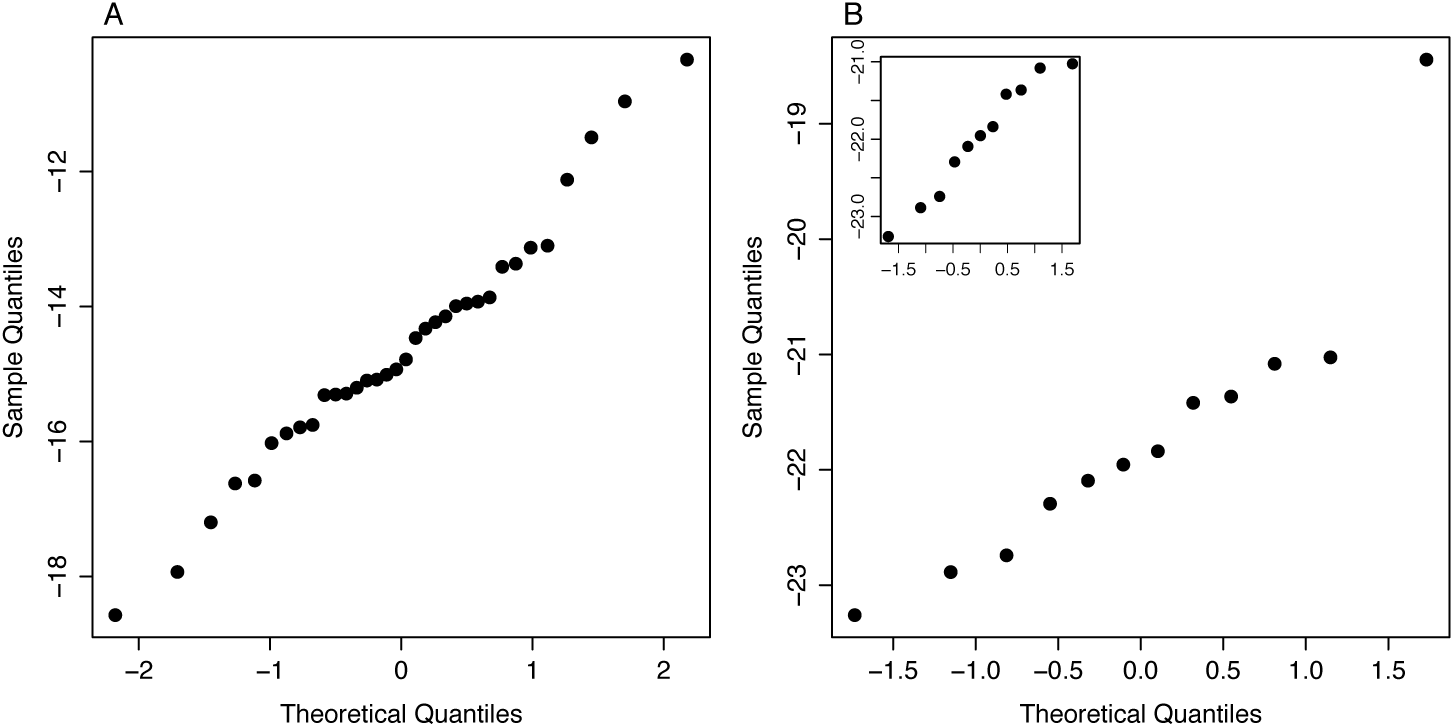
Normal Q-Q plots for the log of (A) accumulation and (B) mutation rate data. The main plot in B includes all twelve mutation rate estimates and the insert excludes *Mesoplasma florum* estimate.

If the accumulation and mutation rate data are log-normally distributed then the distribution of DT is also log-normally distributed with a mean of log_e_(8760) + *m*_*g*_ – *m*_*y*_ and a variance of *v*_*g*_ + *v*_*y*_ – 2Cov(*g,y*), where 8760 is the number of hours per year and *m*_*g*_, *m*_*y*_, *v*_*g*_ and *v*_*y*_ are the mean and variance of the lognormal distributions fitted to the mutation (subscript *g)* and accumulation (subscript *y*) rates. *Cov*(*g,y*) is the covariance between the accumulation and mutation rates. We might expect that species with higher mutation rates also have higher accumulation rates, because the accumulation rate is expected to depend on the mutation rate, but the correlation between the two will depend upon how variable the DT and other factors, such as the strength of selection, are between bacteria. The observed correlation between the log accumulation rate and log mutation rate is negative at -0.47, but there are only four data points, so the confidence intervals on this estimate encompass almost all possible values (-0.99 to 0.89). We explore different levels of the correlation between the accumulation and mutation rates; it should be noted that *Cov*(*g,y*) can be expressed as Sqrt(*v*_*g*_ *v*_*y*_) *Corr*(*g,y*) where *Corr*(*g,y*) is the correlation between the two variables.

The distribution of DTs in the wild inferred using our method is shown in Figure 2. We infer the median doubling time to be 7.4 hours, but there is considerable spread around this even when the accumulation and mutation rates are strongly correlated (Figure 2A); as the correlation increases so the variance in DTs decreases, but the median remains unaffected. The analysis suggests that most bacteria have DTs of between 1 and 100 hours but there are substantial numbers with DTs beyond these limits. For example, even if we assume that the correlation between the accumulation and mutation rate is 0.5 we infer that 10% of bacteria have a DT of faster than one hour in the wild and 4.8% have a DT slower than 100 hours in the wild. If we remove the *Mesoplasma florum* mutation rate estimate from the analysis the median doubling is slightly lower at 5.5 hours, but there is almost as much variation as when this bacterium is included; at a correlation is 0.5 we infer that 13.6% of bacteria have a DT faster than one hour in the wild and 3% have a DT slower than 100 hours.

**Figure 2.**
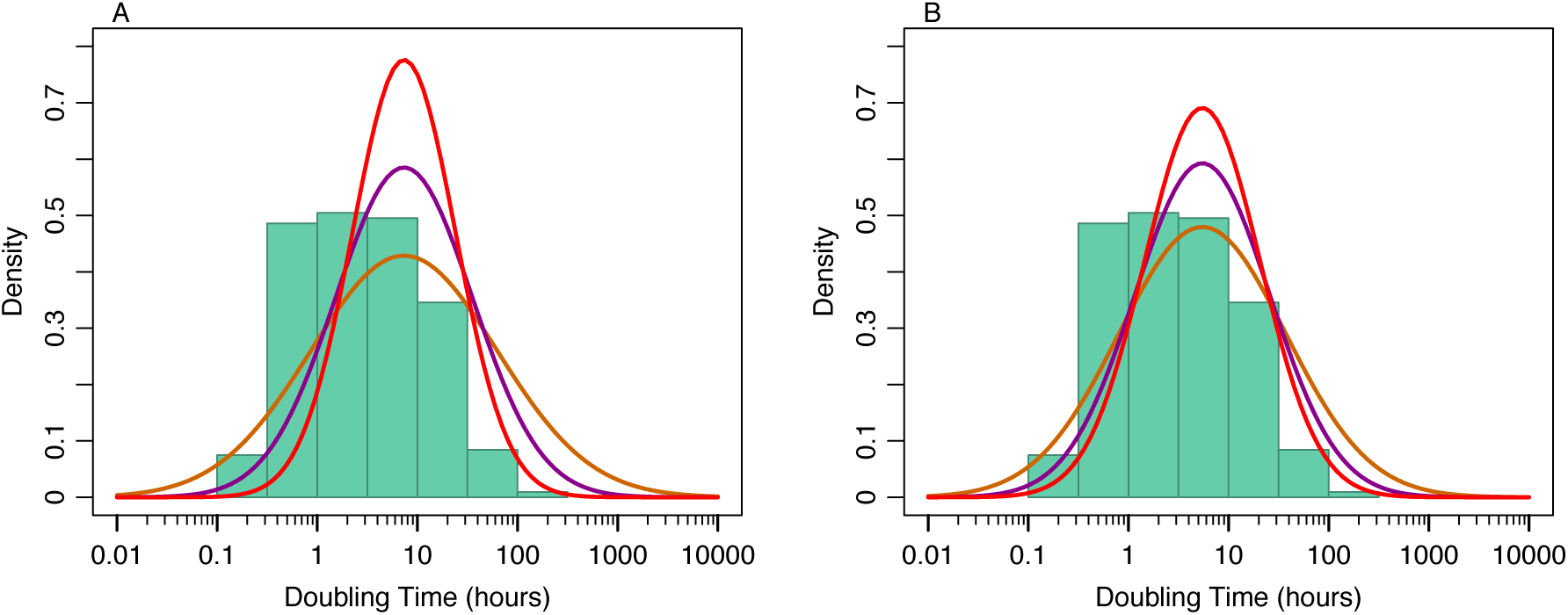
The distribution of DTs amongst bacteria inferred assuming different levels of correlation between the accumulation and mutation rates - orange r = 0, purple r = 0.5 and red r = 0.75. We also show the distribution of lab DTs (green histogram) from a compilation of over 200 species made by Vieira-Silva and Rocha (2010). In panel A we include all mutation rate estimates and in panel B we exclude the mutation rate estimate for *Mesoplasma florum*.

To investigate how robust these conclusions are to statistical sampling, we bootstrapped the accumulation and mutation rate estimates, refit the log-normal distributions and reinferred the distribution of DT. The 95% confidence intervals for the median are quite broad at 3.1 to 19 hours (2.6 to 11.2 hours if we exclude *M. florum*). However, all bootstrapped distributions show substantial variation in the DT with a substantial fraction of bacteria with long DTs and also some with very short DTs (Figure 3).

**Figure 3.**
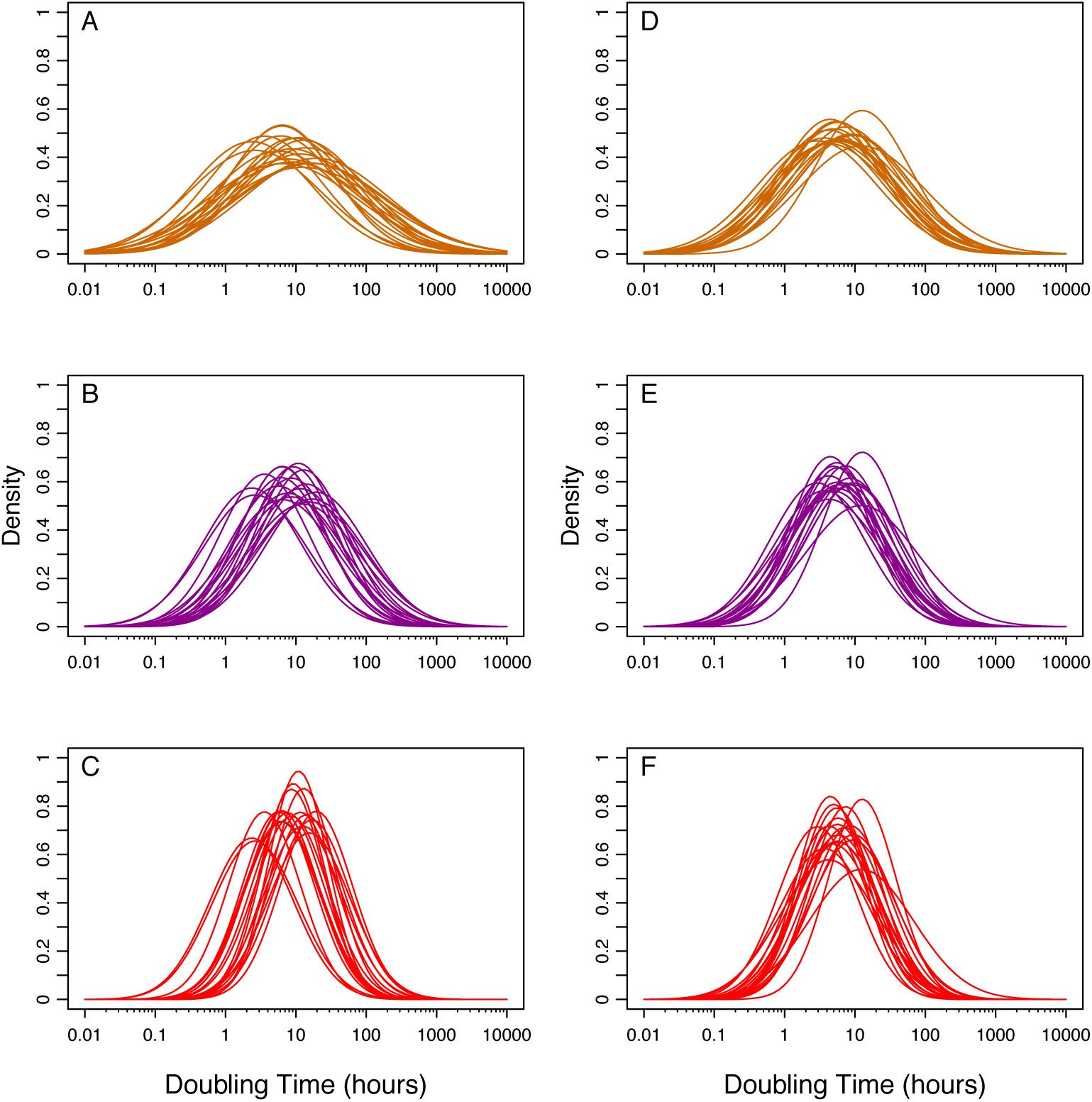
DT distributions inferred by bootstrapping the accumulation and mutation rate data and refitting the log-normal distributions to both datasets. Each plot shows 20 bootstrap DT distributions assuming different levels of correlation between the accumulation and mutation rates - orange r = 0, purple r = 0.5 and red r = 0.75. A, B and C include all mutation rate estimates and D,E, and F show the analysis after removal of the *Mesoplasma florum* mutation rate estimate.

It is of interest to compare the distribution of DTs in the wild to the distribution of lab DTs (Figure 1). The distributions are different in two respects. First, the median lab DT of 3 hours is less than half the median wild DT of 7.4 hours (5.5 hours without *M. florum*); the two are significantly different (p = 0.025 with *M. florum* and p = 0.042 without *M.florum*, inferred by bootstrapping each dataset and recalculating the medians). Second, many more bacteria are inferred to have long DTs in the wild than in the lab.

## Discussion

The DT of most bacteria in their natural environment is not known. We have used estimates of the rate at which bacteria accumulate mutations in their natural environment and estimates of the rate at which they mutate in the laboratory to estimate the DT for several bacteria and infer the distribution of DTs across bacteria. We estimate that DT are generally longer in the wild than in the lab, but critically we also infer that DTs vary by several orders of magnitude between bacterial species and that many bacteria have very slow DTs in their natural environment.

The method, by which we have inferred the DT in the wild, makes three important assumptions. We assume that the mutation rate per generation is the same in the lab and in the wild. However, it seems likely that bacteria in the wild will have a higher mutation rate per generation than those in the lab for two reasons. First, bacteria in the wild are likely to be stressed and this can be expected to elevate the mutation rate (20–24). Second, if we assume that DTs are longer in the wild than the lab then we expect the mutation rate per generation to be higher in the wild than in the lab because some mutational processes do not depend upon DNA replication. The relative contribution of replication dependent and independent mutational mechanisms to the overall mutation rate is unknown. Rates of substitution are higher in firmicutes that do not undergo sporulation suggesting that replication is a source of mutations in this group of bacteria (25). However, rates of mutation accumulation seem to be similar in latent versus active infections of *M. tuberculosis*, suggesting that replication independent mutations might dominate in this bacterium (26, 27).

The second major assumption is that the rate at which mutations accumulate in the wild is equal to the mutation rate per year; in effect, we are assuming that all mutations are effectively neutral, at least over the timeframe in which they are assayed (or that some are inviable, but the same proportion are inviable in the wild and the lab). In those accumulation rate studies, in which they have been studied separately, non-synonymous mutations accumulate more slowly than synonymous mutations; relative rates vary from 0.13 to 0.8, with a mean of 0.57 (Table S3). There is no correlation between the time-frame over which the estimate was made and the ratio of non-synonymous and synonymous accumulation rates (r = 0.2, p = 0.53). We did not attempt to control for selection because the relative rates of synonymous and non-synonymous accumulation are only available for a few species, and the relative rates vary between species. However, we can estimate the degree to which more selection against mildly deleterious non-synonymous accumulations in the wild causes the DT to be underestimated as follows. The observed rate at which mutations accumulate in a bacterial lineage is *μ*_*obs*_ = *α μ*_*true*_ *δ*_*i*_ + (1-*α*)(1-*β*) *μ*_*true*_ *δ*_*s*_ + (1-*α*) *β μ*_*true*_ *δ*_*n*_, where *α* is the proportion of the genome that is non-coding and *β* is the proportion of mutations in protein coding sequence that are non-synonymous. *δ*_*x*_ is the proportion of mutations of class *x* (*i* = intergenic, *s* = synonymous, *n* = non-synonymous) that are effectively neutral. *α* and *β* are approximately 0.15 and 0.7 in our dataset. Although there is selection on synonymous codon use in many bacteria (28), selection appears to be weak (29) we therefore assume that *δ*_*s*_ = 1. This implies, from the rate at which non-synonymous mutations accumulate relative to synonymous mutations, that *δ*_*n*_ = 0.6. A recent analysis of intergenic regions in several species of bacteria has concluded that selection is weaker in intergenic regions than at non-synonymous sites, we therefore assume that *δ*_*i*_ = 0.8 (30). Using these estimates suggests that selection leads us to an underestimate of the true mutation rate per year in the wild by ~27%; this in turn means we have underestimated the DT by ~27%, a relatively small effect.

We have also assumed that there is no phylogenetic inertia in the accumulation and mutation rates; i.e. that closely related species do not have more similar accumulation and mutations rates, than more distantly related species. If there is phylogenetic inertia in one of the variables but not the other then our estimate of the distribution of DTs only applies to the taxa that have been sampled, and if there is phylogenetic inertia in both, then we would have to take account of the phylogenetic structure in our analysis. We tested for a phylogenetic signal by constructing a tree using 16S rRNA sequences and then applying the tests of Pagel (1999) (31) and Blomberg et al. (2003) (32). The accumulation rate estimates show significant signal using Pagel’s test (λ = 0.68, *p* = 0.001) but not Blomberg et al. (*K* = 0.0005, *p* = 0.37). The mutation rate estimates show no significant phylogenetic signal with either test (Pagel: λ = 0.79 *p* = 0.24; Blomberg et al.: *K* = 0.58, *p* = 0.2). It therefore seems that there is not a strong phylogenetic signal and our method should be satisfactory.

Despite the assumptions we have made in our method, our estimate of the DT of *P. aruginosa* of 2.3 hours in a cystic fibrosis patient is very similar to that independently estimated using the ribosomal content of cells of between 1.9 and 2.4 hours (3). There is also independent evidence that there are some bacteria that divide slowly in their natural environment. The aphid symbiont *Buchnera aphidicola* is estimated to double every 175-292 hours in its host (33, 34), and *Mycobacterium leprae* doubles every 300-600 hours on mouse footpads (35–37), not its natural environment, but one that is probably similar to the human skin. Furthermore, in a recent selection experiment, Avrani et al. (2017) (38) found that several *E. coli* populations, which were starved of resources, accumulated mutations in the core RNA polymerase gene. These mutations caused these strains to divide more slowly than unmutated strains when resources were plentiful. Interestingly these same mutations are found at high frequency in unculturable bacteria, suggesting that there is a class of slow growing bacteria in the environment that are adapted to starvation.

Korem et al. (2015) (39) have recently proposed a general method by which the DT can be potentially estimated. They note that actively replicating bacterial cells have two or more copies of the chromosome near the origin of replication but only one copy near the terminus, if cell division occurs rapidly after the completion of DNA replication. Using next generation sequencing, they show that it is possible to assay this signal and that the ratio of sequencing depth near the origin and terminus is correlated to bacterial growth rates *in vivo*. Brown et al. (2016) (40) have extended the method to bacteria without a reference genome and/or those without a known origin and terminus of replication. In principle, these measures of cells performing DNA replication could be used to estimate the DT of bacteria in the wild. However, it’s unclear how or whether the methods can be calibrated. Both Korem et al. (2015) and Brown et al. (2016) find that their replication measures have a median of ~1.3 across bacteria in the human gut. However, a value of 1.3 translates into different relative and absolute values of the DT in the two studies. Brown et al. (2016) show that their measure of replication, iRep, is highly correlated to Korem et al.’s (2015) measure, PTR, for data from *Lactobacillus gasseri*; the equation relating the two statistics is iRep = -0.75 + 2 PTR. Hence, when PTR = 1.3, iRep = 1.85 and when iRep = 1.3, PTR = 1.03. The two methods are not consistent. They also yield very different estimates for the absolute DT. Korem et al. (2015) show that PTR is highly correlated to the growth rate of *E. coli* grown in a chemostat. If we assume that the relationship between PTR and growth rate is the same across bacteria *in vivo* and *in vitro*, then this implies that the median DT for the human microbiome is ~2.5 hours. In contrast, Brown et al. (2016) estimate the growth rate of *Klebsiella oxytoca* to be 19.7 hours in a new-born baby using faecal counts and find that this population has an iRep value of ~1.77. This value is greater than the vast majority of bacteria in the human microbiome and bacteria in the Candidate Phyla Radiation, suggesting that most bacteria in these two communities replicate very slowly. These discrepancies between the two methods suggest that it may not be easy to calibrate the PTR and iRep methods to yield estimates of the DT across bacteria.

In summary, the availability of accumulation and mutation rate estimates allows us to infer the DT for bacteria in the wild, and the distribution of wild DTs across bacterial species. These DT estimates are likely to be underestimates because the mutation rate per generation is expected to be higher in the wild than in the lab, and some mutations are not generated by DNA replication. Our analysis therefore suggests that DTs in the wild are typically longer than those in the lab, that they vary considerably between bacterial species and that a substantial proportion of species have very long DTs in the wild.

## Materials and methods

We compiled estimates of the accumulation and mutation rate of bacteria from the literature. We only used mutation rate estimates that came from a mutation accumulation experiment with whole genome sequencing. If we had multiple estimates of the mutation rate we summed the number of mutations across the mutation accumulation experiments and divided this by the product of the genome size and the number of generations that were assayed. We averaged the accumulation rate estimates where we had multiple estimates from the same species. We recalculated the accumulation rates in two cases in which the number of accumulated mutations had been divided by an incorrect number of years: *E. coli* (5) and *Helicobacter pylori* (41). For *E. coli*, we reestimated the accumulation rate using BEAST by constructing sequences of the SNPs reported in the paper and the isolation dates. For, *Helicobacter pylori* the 3-year and 16-year strains appear to form a clade to the exclusion of the 0-year strain because they share some differences from the 0-year strain. If the number of substitutions that have accumulated between the common ancestor of the 3-year and 16-year strain and each of the two strains are *S*_*3*_ and *S*_*16*_ respectively then the rate of accumulation can be estimated as (*S*_*16*_-*S*_*3*_)/(13 years x genome size). For the isolates NQ1707 and NQ4060 we have estimated the accumulation rate to be 5 × 10^−6^ and for NQ1671 and NQ4191 5.9 × 10^−6^. We excluded some accumulation rate estimates for a variety of reasons. We only considered accumulation rates sampled over an historical timeframe of at most 1500 years. Most of our estimates of the accumulation rate are for all sites in the genome, so we excluded cases in which only the synonymous accumulation rate was given. We also excluded accumulation rates from hypermutable strains. Accumulation and mutation rate estimates used in the analysis are given in supplementary tables S1 and S2 respectively.

The estimate of the standard error associated with our estimate of the doubling time was obtained using the standard formula for the variance of a ratio: V(*x/y*) = (M(*x*)/M(*y*))^2^(V(*x*)/M(*x*)^2^+V(*y*)/M(*y*)^2^) where M and V are the mean and variance of *x* and *y*. The variance for the accumulation rate was either the variance between multiple estimates of the accumulation rate if they were available, or the variance associated with the estimate if there was only a single estimate. The variance associated with the mutation rate was derived by assuming that the number of mutations was Poisson distributed.

We fit log-normal distributions to the accumulation and mutation rate data by taking the log_e_ of the values and then fitting a normal distribution by maximum likelihood using the *FindDistributionParameters* in *Mathematica*. Normal Q-Q plots for the accumulation and mutation rate data were produced using the qqnorm function in R version 1.0.143. In fitting these distributions, we have not taken into account the sampling error associated with the accumulation and mutation rate estimates. However, these sampling errors are small compared to the variance between species: for the accumulation rates the variance between species is 3.9 × 10^−11^ and the average error variance is an order of magnitude smaller at 3.6 × 10^−12^; for the mutation rate data, the variance between species is 7.5 × 10^−18^ and the average variance associated with sampling is more than two orders of magnitude smaller at 1.8 × 10^−20^. Note that we cannot perform these comparisons of variances on a log-scale because we do not have variance estimates for the log accumulation and mutation rates.

To estimate phylogenetic signal in the accumulation and mutation rates we generated phylogenetic trees for each set of species in the two datasets. 16S rRNA sequences were downloaded from the NCBI genome database (https://www.ncbi.nlm.nih.gov/genome/) and aligned using MUSCLE (Edgar 2004) performed in Geneious version 10.0.9 (http://www.geneious.com, Kearse et al. 2012). From these alignments, maximum likelihood trees were constructed in RAxML (43) and integrated into the tests of Pagel (1999) (31) and Blomberg et al. (2003) (32) to the log_10_(accumulation rates) and log_10_ (mutation rates) implemented in the phylosig function in the R package phytools v.0.6 (44).

## Acknowledgements

We are very grateful to Michael Lynch for sharing his mutation rate estimates prior to publication. DJW is a Sir Henry Dale Fellow, jointly funded by the Wellcome Trust and the Royal Society (grant no. 101237/Z/13/Z).

